# Structural Basis of Aquaporin-4 Autoantibody Binding in Neuromyelitis Optica

**DOI:** 10.1101/2024.05.12.592631

**Authors:** Meghna Gupta, Nitesh Kumar Khandelwal, Andrew Nelson, Peter Hwang, Sergei Pourmal, Jeffrey L. Bennett, Robert M. Stroud

## Abstract

Neuromyelitis Optica (NMO) is an autoimmune disease of the central nervous system where pathogenic autoantibodies target the human astrocyte water channel aquaporin-4 causing neurological impairment. Autoantibody binding leads to complement dependent and complement independent cytotoxicity, ultimately resulting in astrocyte death, demyelination, and neuronal loss. Aquaporin-4 assembles in astrocyte plasma membranes as symmetric tetramers or as arrays of tetramers. We report molecular structures of aquaporin-4 alone and bound to Fab fragments from patient-derived NMO autoantibodies using cryogenic electron microscopy. Each antibody binds to epitopes comprised of three extracellular loops of aquaporin-4 with contributions from multiple molecules in the assembly. The structures distinguish between antibodies that bind to the tetrameric form of aquaporin-4, and those targeting higher order orthogonal arrays of tetramers that provide more diverse bridging epitopes.

**One-Sentence Summary:** Pathogenic autoantibodies in Neuromyelitis Optica distinguish multiply presented epitopes on arrays of human aquaporin-4.

---

Neuromyelitis Optica (NMO) is a severe and debilitating autoimmune disease of the central nervous system (CNS). In most cases (≥80%), NMO is mediated by antibodies that target the extracellular loops of the human water channel aquaporin-4 (AQP4) in the plasma membrane (*1*). Pathogenic AQP4 immunoglobulin G autoantibodies (AQP4-IgGs), when they interact with AQP4 on astrocytes of the CNS, initiate tissue injury and cause neurological impairment through both lytic and non-lytic mechanisms (*2*). AQP4-IgG mediated effector functions, complement-mediated cytotoxicity (CDC), and antibody dependent cell-mediated cytotoxicity (ADCC) cause targeted astrocyte lysis, promote immune cell infiltration, and drive demyelination, axonal injury, and neuronal destruction (*3-7*). CDC activation is due to binding of several IgGs into a multimeric platform for C1q assembly (16). Current therapies are based on strategies that may result in significant short- and long-term adverse events (*8*). To advance NMO therapy and diagnosis, we sought to define the structural basis of autoantibody binding, the initial event of the disease, using patient-derived AQP4-specific recombinant antibodies (rAbs).

The epitopes in NMO are formed by three well-ordered extracellular loops (loops A, C, E) displayed in each AQP4 molecule. Four monomeric water channels form symmetric tetramers, as in all aquaporins, therefore the conformational epitopes can include extracellular loops from the four adjacent monomers. A higher order assembly of AQP4 is formed by a splice variant with a start site at Met23 (M23). M23 lacks the first 22 amino acids as compared to the full-length AQP4 (M1) on the cytoplasmic side of the plasma membrane. M23 preferentially forms two-dimensional (2D) <u>O</u>rthogonal <u>A</u>rrays of tetrameric AQP4 <u>P</u>articles (OAPs), and displays the identical extracellular epitopes as the tetramers. M1 disfavors OAP formation, in part due to palmitoylation at Cys13 and Cys17 on the cytoplasmic side. Therefore, OAPs present additional copies of the epitopes comprised of A, C, E loops arrayed different orientations by adjacent tetramers within the OAPs. The distribution of M1 to M23 is determined by their relative expression and a combination of post-translational modifications (*9*). Higher proportions of M1 limit the size of OAPs suggesting that M1 is incorporated into the lattice and limits its extent (*9*).

Patient-derived anti-AQP4 monoclonal recombinant antibodies (AQP4 rAbs) do not recognize linear peptide epitopes that correspond to the loops A, C, or E, indicating that combinations of the folded three-dimensional (3D) structures of these loops displayed on the external surface of AQP4 are the basis of the conformational epitopes (*10, 11*). In cell-based assays, a subset of purified AQP4-IgGs bind equally well to M1 as to M23 AQP4 indicating that their epitopes are contained within the AQP4 tetramer. However, most AQP4 rAbs have higher affinities for M23 OAPs. These factors imply that most AQP4 rAbs bind to conformational epitopes comprised of multiple interactions from neighboring tetramers. The comparison of Fab fragment and divalent IgG binding in these assays did not show significant difference in their binding affinities, suggesting that each IgG binds the target with one Fab fragment, without any avidity effect of bivalent binding by IgGs (*12*).

Mutagenesis and changing M1:M23 expression ratios demonstrate that the higher affinity of Fab binding is due to OAP formation. Here we determined the structural basis of binding of human AQP4 to the patient-derived AQP4-rAb Fab fragments from-(a) rAB58, that binds equally well to both M1 and M23 AQP4 isoforms and (b) rAB186, that binds ∼55 times better (lower K_d_) to the M23 OAPs than to M1 (*12*).

## CryoEM structure of human AQP4 in lipid nanodiscs

The AQP4 M1 isoform was purified as a homo-tetrameric complex and reconstituted into lipid nanodiscs comprised of soybean polar lipid extract surrounded by the membrane scaffold protein MSP-1E3D1 (Fig S1). We determined structure of the AQP4 M1 tetramer to 2.1Å resolution by cryoEM (Fig 1, S2, S3). The water channel and the continuous line of hydrogen bonded water molecules through it are essentially identical to our previously determined AQP4 crystal structure using X-ray diffraction (*13*). To obtain that crystal structure of AQP4 tetramers, AQP4 was treated with trypsin to remove flexible regions of the protein, that then gave a 1.8Å resolution electron density map. For the structure reported here, the full-length protein was reconstituted into lipid nanodiscs to maintain a more native environment and was not treated with trypsin. The first 31 amino acid residues and amino acid residues 69 residues at the C-terminal (254-323) are not seen in the cryoEM map, consistent with the flexibility of these cytoplasmic regions.

**Figure 1.**
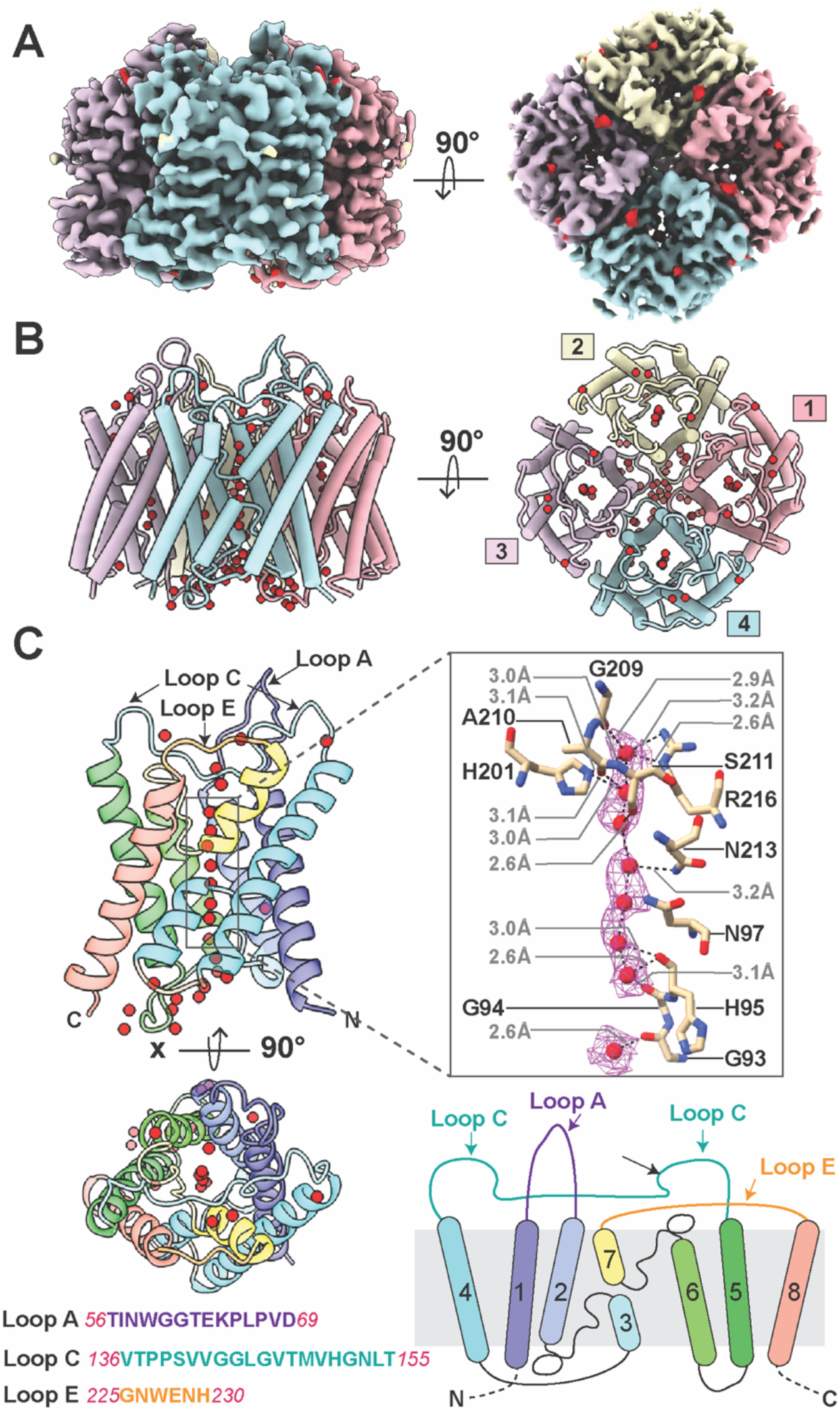
Structure of human AQP4 tetramer in nanodiscs. **(A)** AQP4 density map determined by cryoEM at 2.1 Å resolution, side view and top view showing the extracellular surface of the same. Each monomer is colored differently, blue, pink, lemon, thistle anticlockwise. The four individual water channels contain water molecules (shown in red). **(B)** Side view and top view cartoon of the AQP4 tetramer model built using the density map in (A). The monomers are colored as (A) and numbered anticlockwise **(C)** AQP4 monomer shown in rainbow (from N to C terminus) in the side and top views representing location of water molecules. The three extracellular loops A, C, E are shown in the side view. Key amino acid residues interacting with the waters are highlighted and experimentally determined waters in the density maps are shown with mesh surface. A cartoon of AQP4 monomer depicting the relative organization of transmembrane helices in rainbow. The sequence of each extracellular loop shown.

The structure shows the three extracellular loops to which the AQP4-IgGs bind (*10, 11, 14*). Loop A (56-TINWGGTEKPLPVD-69) between transmembrane helices TM1 and TM2 has a well-defined stable structure and extends the farthest above the plane of the membrane. This loop presents the polar charged side chains of E63, K64 at the top (Fig 1). Loop A is closest to the 4-fold symmetry axis of the tetrameric assembly. Loop C (136-VTPPSVVGGLGVTMVHGNLT-155) between TM4 and TM5 is the longest loop of 20 amino acids, and traverses the width of AQP4. Side chains in the center of the loop C (145-LGVT-148) touch the membrane surface such that there are two distinct epitopes of loop C that are presented; the proximal C loop (136-VTPPSVVGG-144), and the distal C loop epitope (149-MVHGNLT-155). Loop E (225-GNWENH-230) between TM7 and TM8 is the shortest loop, is close to the membrane surface, and is the most distant radially from the 4-fold axis of AQP4.

Densities define the line of hydrogen bonded water molecules throughout the pore of each channel (Fig 1). Water molecules are associated with His201 and Arg216 in the ‘selectivity filter’ of the channel, and at the central water-polarizing site formed by hydrogen bond donors from two Asn residues of the twinned signature sequences of all aquaporins that consist of two -Asn-Pro-Ala-(-NPA-) motifs, one from each half of the sequence. These polarize the central water that positions all waters to orient donor hydrogen bonds toward each side preventing any ion, even hydronium from passing through the channel. The 4-fold symmetry axis in the AQP4 tetramer is surrounded by hydrophobic side chains and is not functional for transport.

Using single amino acid substitutions in loops A, C, and E, and AQP4 rAbs (*5*), two major binding patterns of rAbs were observed: pattern 1 in which AQP4 rAbs mutations of individual residues to alanine in extracellular loops C and E show reduced rAb binding, and pattern 2 AQP4 rAbs where mutations in any of the three loops reduced rAb binding (12). Independent of the pattern type, some AQP4 rAbs are dependent on residues H151-L154 in distal loop C of AQP4 and demonstrate enhanced complement-mediated cytotoxicity (*15*). Binding of two pattern 2 AQP4 rAbs, rAb58, and rAb186, to purified AQP4 in nanodiscs was confirmed using Bio-Layer Interferometry (BLI), and their structures and binding interactions were determined by cryoEM.

### Fab58 spans adjacent monomers in the AQP4 tetramer

2D class averages of cryoEM images demonstrate that each AQP4 tetramer is bound by only a single Fab58 (Fig 2, S4). The 3D structure obtained at 2.5Å resolution shows that one Fab58 binds a compound epitope between two adjacent monomers within a single tetramer (Fig 2, Fig S4, S5). The binding of one Fab58 sterically prevents binding of other Fab58s to other monomers in the tetramer (Fig 4A, 4C). rAb58 binds equally well to tetramers as to OAPs in a cell-based assay with K_d_ values reported between ∼60-150 nM (*15, 16*). There is no structural impediment to water access to the channel entrance from the presence of bound Fab58 (Fig 2).

**Figure 2.**
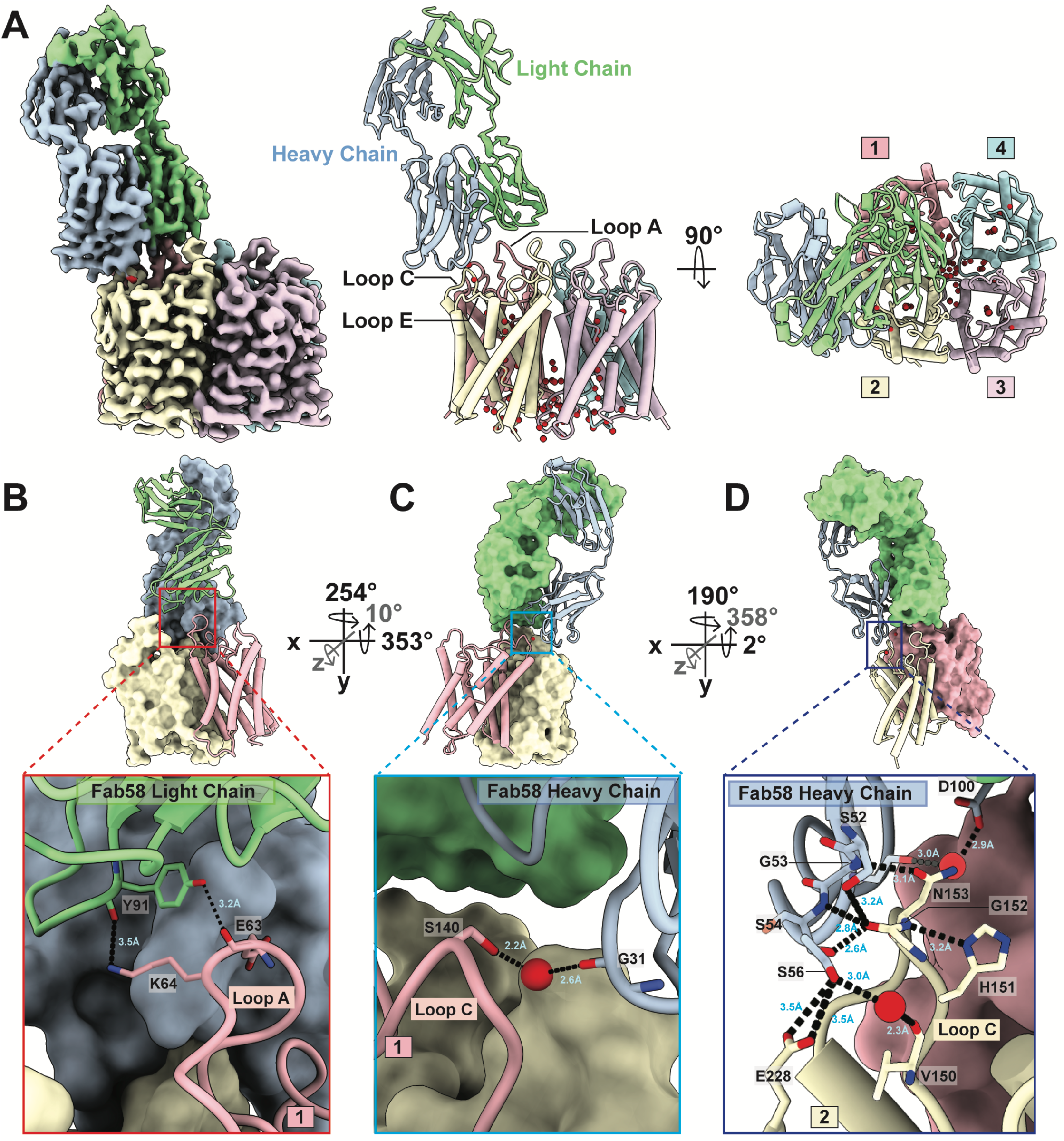
Fab58 binding to human AQP4 M1 tetramer. **(A)** Side view of the cryoEM density map at 2.5 Å resolution showing AQP4 tetramer and Fab58 interacting with the extracellular loops. Fab58 HC is shown in light steel blue and LC in pale green. A cartoon of the side view and top view of the model built using the density map with the same color scheme and numbered anticlockwise. One molecule of Fab58 makes interactions with two neighboring AQP4 monomers in the tetramer. **(B)** Molecular interactions between the LC of Fab58 and loop A of monomer 1 of AQP4 tetramer. Bond length in Å is mentioned in blue and the bonds are shown with a black dotted line. **(C)** Molecular interactions between the HC of Fab58 and loop C of monomer 1 of AQP4 tetramer mediated by a water molecule. **(D)** Molecular interactions between the HC of Fab58 and AQP4 loop C of monomer 2 of AQP4 tetramer. Two water molecules in red play an important role in this interaction.

Fab58 contacts residues of loop A and the proximal end of loop C from one AQP4 monomer and the distal end of loop C from its neighboring monomer as seen anticlockwise from the external side of the cell in the tetramer (Fig 2). Y91 of Fab58 light chain (LC) binds amino acids 63-EK-64 of loop A. while the carbonyl of G31 of Fab58 heavy chain (HC) interacts with loop C amino acid S140 via a water molecule (Fig 2B, 2C). Fab58 HC provides a polar environment and makes hydrogen bonds from the N-H of G53 and D100 HC via water to N153, and three hydrogen bonds to the carbonyl of G152 of distal loop C from the adjacent monomer. S56 HC makes a hydrogen bond via a water molecule to the carbonyl of V150, and to the carboxyl group of E228 (E loop) (Fig 2D).

The side chains of two aromatic rings W32 in LC and Y101 in HC of Fab58 sandwich the aliphatic K64 side chain in the ‘-EKP-’ segment of loop A. These residues on the Fab are each buttressed by other interactions within the Fab58 R30 in LC and D100 in HC suggesting that this pi-aliphatic cation-pi interaction can be very favorable for antigen binding. The terminal NH_3_^+^ of K64 is hydrogen bonded to the carbonyl of Y91, interactions that together can serve to fix the aliphatic chain of K64 and can then account for a large component of the association energy of Fab58 to AQP4 (Fig S6).

Substitutions of certain residues in the extracellular loops of AQP4 to alanine diminish rAb58 binding as measured by fluorescence-based imaging, supporting a role in target sites bound by Fab58 (*11*). N153A diminished binding by both pattern 1 and 2 Fabs while N153Q only diminished pattern 2 rAbs (*11*). N153 lies in the interaction between Fab58 and AQP4 (Fig 2D). rAb58 engages all three extracellular loops and is not sensitive to mutagenesis of a H151/L154 in loop C (*15*). Neither H151 nor L154 are close to the interaction interface hence not part of the epitope for Fab58. Mutation of K64A that removes the aliphatic side chain reduces rAb58 binding by ∼80% the largest amount when compared with mutations in nearby residues from T62 to V68 (-TEKPLPV-) (*11*). In summary, rAb58 binds with the similar affinity to M1 tetramers as to M23 OAPs since the compound epitope overlaps loops A and C from two adjacent monomers within a single tetramer of AQP4 (Fig 2A, 4A, 4C) (*12*).

### Fab186 bridges AQP4 tetramers in OAPs

CryoEM images of Fab186-AQP4 show statistical binding of up to four Fabs per AQP4 tetramer (Fig S7, S9). The fourth Fab is visible only at lower contour of the map, probably due to statistical occupation (Fig S9). Each Fab186 bridges two monomers within the tetramer but at an angle away from the 4-fold axis such that the rAb186-AQP4 interface can also incorporate epitopes from neighboring tetramers in the OAPs (Fig 3, 4). And rAb186 binds ∼20-fold tighter to M23 OAPs than to M1 tetramers (*12*). Such inter-tetrameric conformational epitopes provide more diversity than available within the tetramer. N153 is in a central position and the HC of Fab186 makes multiple contacts through the side chain as well as the main chain (Fig 3B). Y54 and Y102 of the Fab186 HC clasp S140 from the proximal part of the loop C from the neighboring AQP4 monomer (Fig 3C).

**Figure 3.**
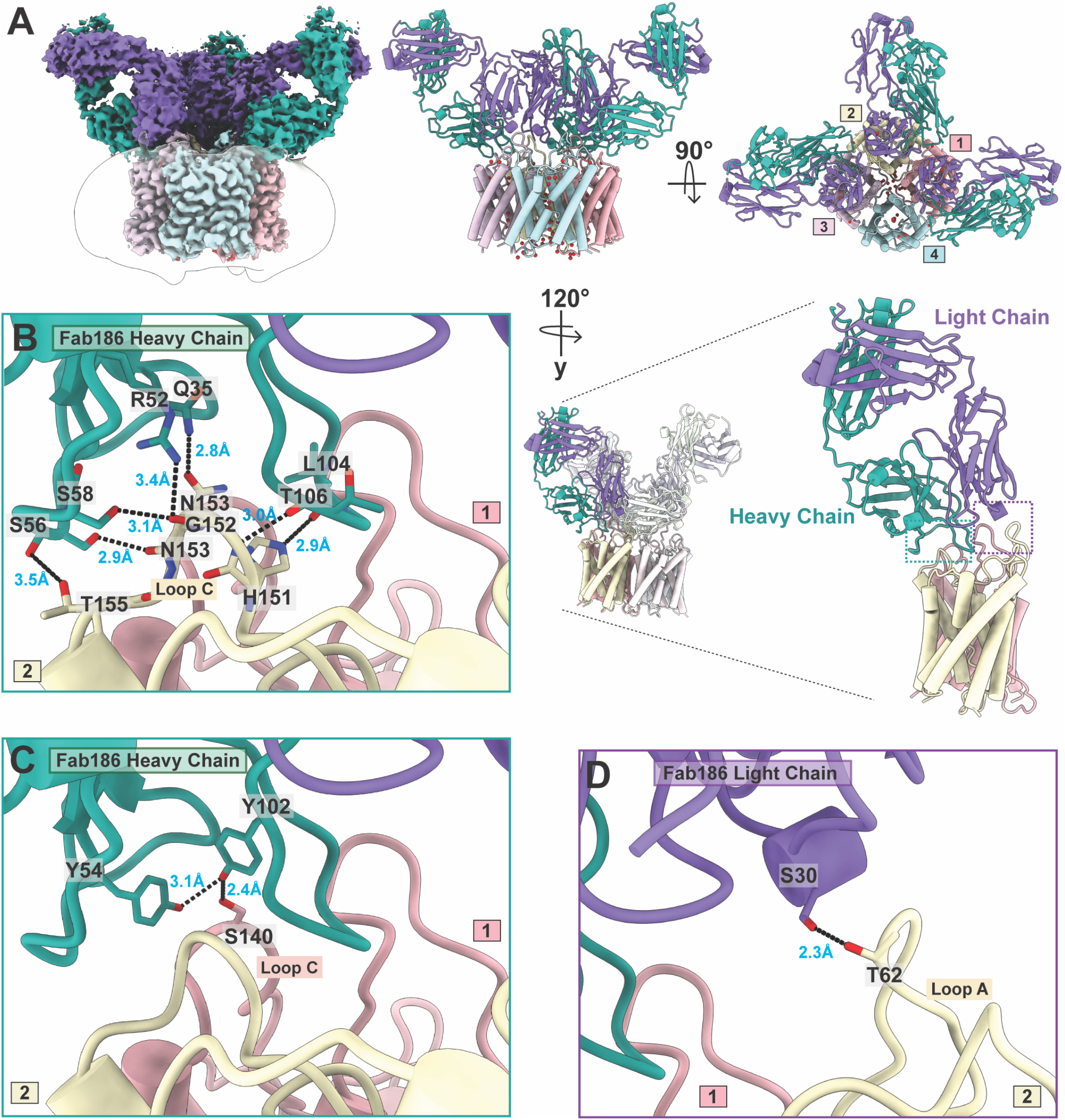
Fab186 binding to human AQP4 M1 tetramer. **(A)** Side view of the cryoEM density map surrounded by a detergent micelle at 2.9 Å resolution showing AQP4 tetramer and three Fab186 molecules interacting with the extracellular loops. Fab186 HC is shown in light sea green and LC in medium purple. A cartoon of the side view and top view of the model built using the density map with the same color scheme and numbered anticlockwise. One Fab186 makes interactions with two neighboring AQP4 monomers in the tetramer. **(B)** Molecular interactions between the HC of Fab186 and loop C of monomer 2 of AQP4 tetramer. Bond length in Å is in blue and hydrogen bonds are shown with black dotted line. **(C)** Hydrogen bonded interactions between the HC of Fab186 and loop C of monomer 1 of AQP4 tetramer. **(D)** Hydrogen bonded interactions between the LC of Fab186 and loop A of monomer 2 of AQP4 tetramer.

Both proximal and distal parts of loop C contribute to this compound epitope from adjacent monomers. S30 of the Fab186 LC hydrogen bonds with T62 of loop A (Fig 3D). There is no interaction with the following 63-EKP-65 though mutations in these residues may change the orientation of the loop and T62 side chain thus causing sensitivity (*11*).

Fab186 has two extended loops one from HC, another from the LC. The extended loop from HC engages the distal loop C H151/L154 of AQP4 explaining the sensitivity of Fab186 to mutations in distal C-loop (Fig 4E) (*15*). L104, T106 of Fab186 HC directly insert into the AQP4 cleft where H151 is presented on AQP4 loop C (Fig 3B, 4E). There is no direct interaction with L154 in our structure, emphasizing that mutations have effects on nearby conformations of the loop.

**Figure 4.**
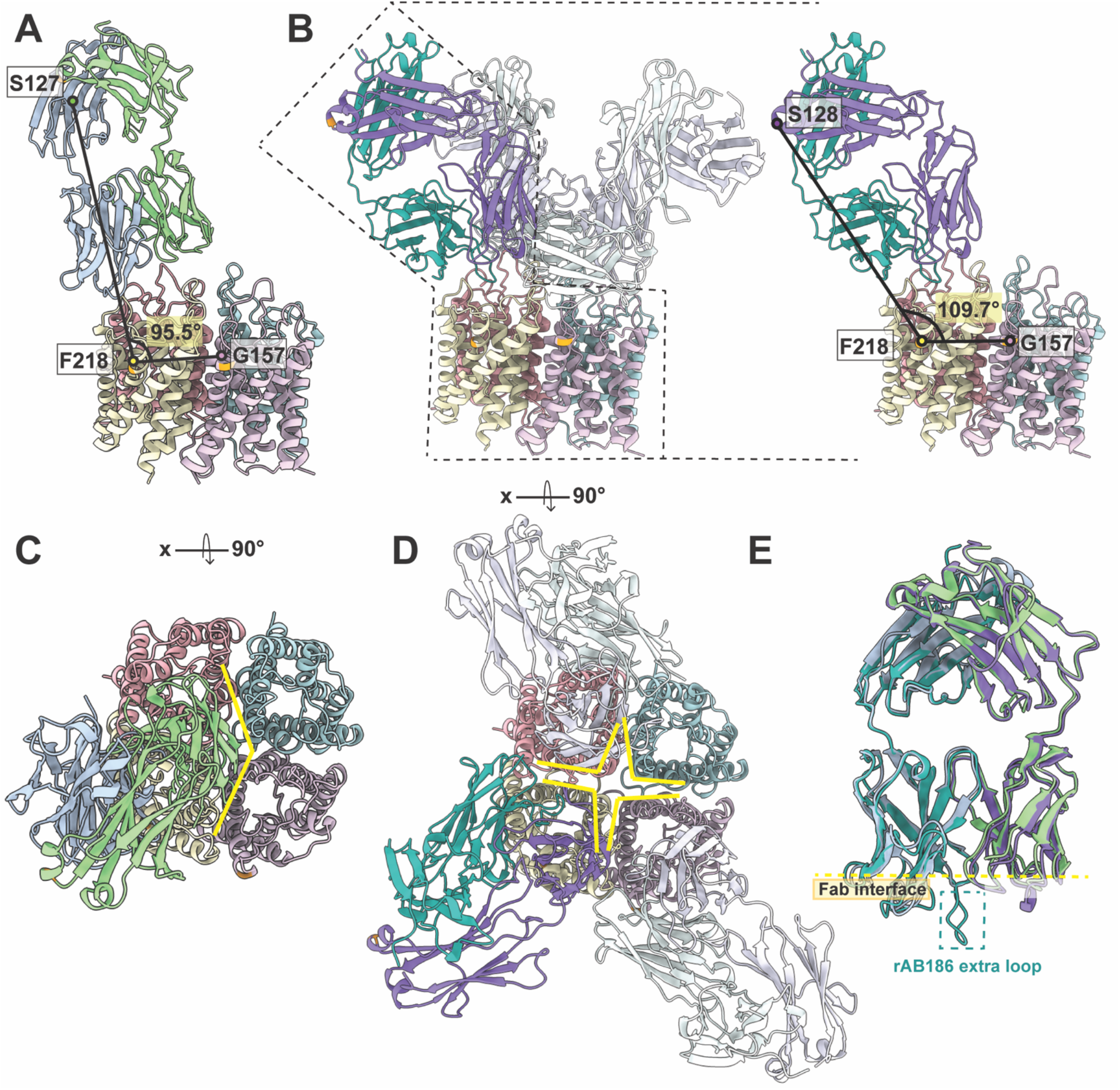
Comparison between Fab58 and Fab186 binding on AQP4 tetramer. **(A)** Fab58 binds on AQP4 surface with an angle of 95.5° between selected labelled Cα atoms while **(B)** Fab186 binds at 109.7° measured using the orthologous amino acid residues to compare Fab58 and Fab186 binding. **(C)** Occupancy of Fab58 on AQP4 tetramer shown in the top view exhibiting steric limitation to accommodate another Fab58 molecule. **(D)** Fab186 binding at a steeper angle enables efficient binding of four Fabs. We see full density for three molecules of Fab while fourth molecule has a weak density as shown in figure S9. **(E)** Overlay of Fab58 and Fab186 structures shows their different loops relative to a constant template. Fab186 has extended regions and one of these take part in key interaction with H151 of loop C differentiating it from Fab58.

The binding angles and orientation of Fab58 and Fab186 are different though both are sensitive to mutations in all three loops of AQP4 (Fig 4). A linear morph between models of Fab58 and Fab186 binding over two AQP4 monomers highlights the differences in interaction (Movie S1).

### Modelling the interaction of rAb186 with OAP arrays

To see which residues of neighboring tetramers in OAPs might interact with rAb186, M23 tetramers of AQP4 in OAPs were modeled based on known structures. In vitro, M23 can form double layers of orthogonal arrays that face each other via their extracellular surfaces as determined using electron diffraction analysis (*17*) (Fig S10). Hence, this interaction between the extracellular leaflets results in compaction of the loops A, C, and E, between the interfaces relative to the structures we determine here. Therefore, to model the cellular surface we replaced each tetramer in a single layer of the crystal lattice by the M1 tetramer structure. (Fig S11). As evidence that the single layer of the double layered lattices is the physiological lattice, the unit cell dimensions of the arrays in rat M23 AQP4 layers a=b=69.0Å are identical to those of the M23 single layer arrays formed in human OAPs in cells (*17*).

The interactions between tetramers in the single layer are also highly conserved in other species where lattice formation is also found, though there is no yet recognized function to these OAP lattices. Using a (‘) to signify an amino acid from an adjacent tetramer in the lattice, specificity of each inter-tetramer close interaction involves W231-L’160/G’157 and W’231-L160/G157, and of Y250 -R’108 and Y’250-R108, and one dimeric pair interaction of I239-I’239 (*17*). G157, W231, and I239 are the hydrophobic resides that constitute the extracellular side of the inter-tetramer interactions. On the cytoplasmic side R108 and Y250 stabilize the interface (*17*). These interactions provide the rationale for the specificity in the tetramer-tetramer interface that drives the in-plane lattice formation. These inter-tetramer residues are all conserved between human, rat, mouse, bovine, with only one conservative change in mouse from I239 to M239 suggesting some yet undefined physiological role. Therefore, this lattice reflects the single layer OAPs that interact with autoimmune rAbs to augment the immune response and allows us to ask how the rAb186 might gain additional interactions from the neighboring tetramers.

The structure of Fab186 is angled with its major axis ∼20° away from perpendicular towards the membrane plane (Fig 4) and forms close contacts of the HC with two monomers of a neighboring tetramer in the model OAP (Fig 5). The observed interaction with the tetramer shows that the Q16 and S85 of HC of Fab186 HC could interact with E63 of loop A, N153 of distal loop C in the neighboring tetramer in the OAP array (Fig 5A, B; Movie S2). These interactions with OAPs can account for rAb186 binding with ∼50-fold higher affinity than to M1 tetramers of AQP4 (*16*). This factor corresponds energetically to ∼3-4.5kcal/M and can be accounted for by these interactions with an adjacent tetramer in OAPs.

**Figure 5.**
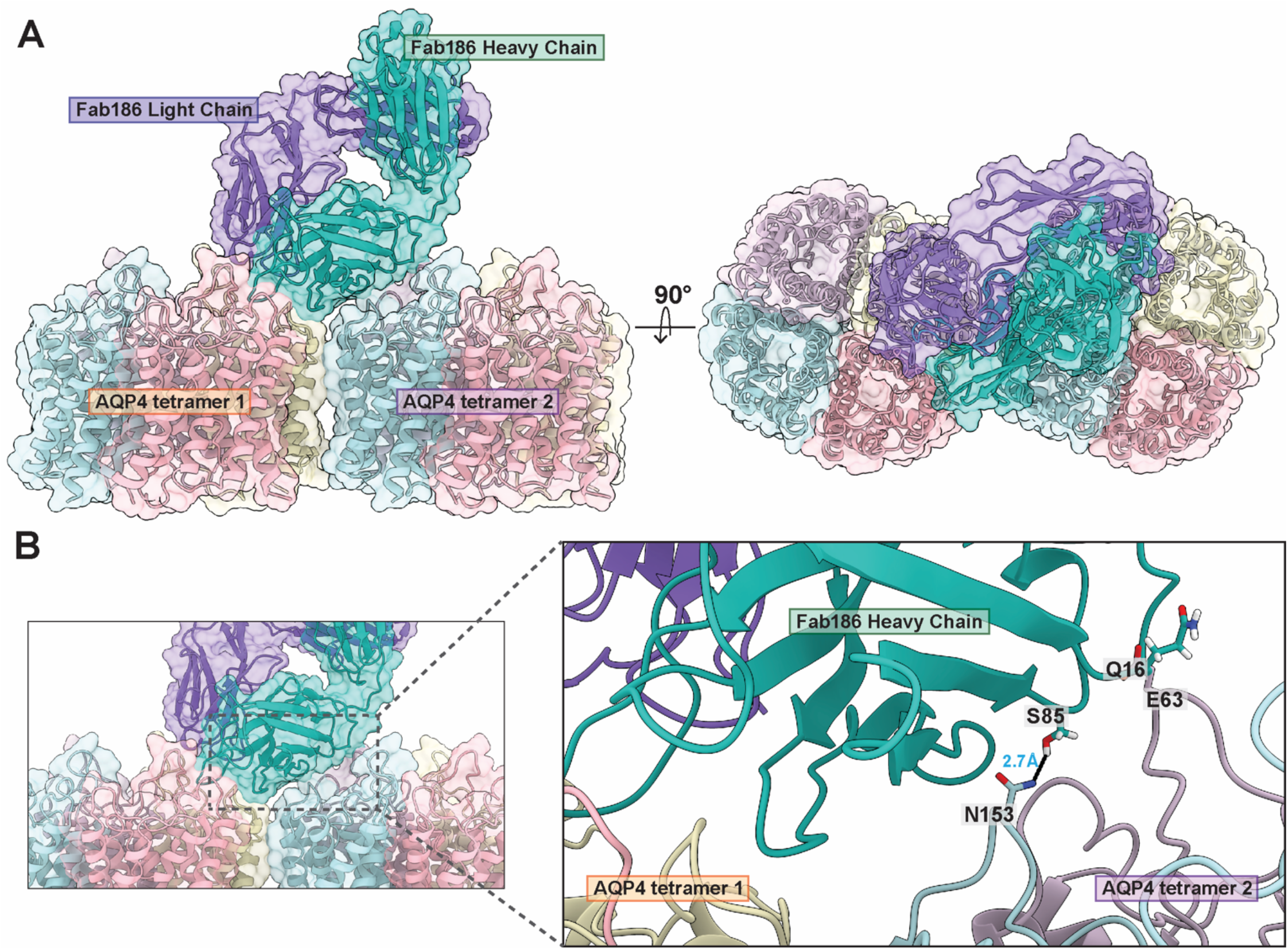
Understanding potential rAB186 association with OAPs. **(A)** Human AQP4 structure superposed on the two neighboring tetramers of rat AQP4 M23 isoform tetramers from 2D crystal lattice structure based on electron diffraction analysis (PDB ID 2D57). Two of the monomers in one of the tetramers were replaced with a minimal Fab186 bound structure consisting of two monomers of AQP4 and one molecule of Fab186. These space filling representations emphasize the proximity of the Fab186 to the neighboring tetramer, favored by AQP4 M23 OAPs. The side view and top view exhibiting the possibility of Fab186 interaction spanning over two AQP4 tetramers. **(B)** Closeup of the Fab186 HC potential interactions (as in A) with loop A and loop C key of the neighboring tetramer.

## Discussion

We determined the structure of M1 AQP4 in lipid nanodiscs versus our previous crystal structure in detergent micelles (*13*). The cryoEM structure is identical to the crystal structure in conformation even though the flexible regions of the protein were removed by proteolysis to form the crystal structure. These same residues are not visible in the cryoEM structure presumably due to their flexibility on the cytoplasmic side. This serves as evidence for the ordered structure of the three extracellular loops that may play into the immune surveillance of structured epitopes.

AQP4 conducts water molecules at close to the diffusion limit for a pore of this size in response to any osmotic pressure as evoked by ion conductance during neuronal action (*18-20*). The water channel contains a continuous single file of water molecules supported by hydrogen bond acceptor carbonyl oxygens along the entire channel pathway (Fig 1). A characteristic structure of twin -NPA-motifs in the center of the human AQP4 channel determine that the central water presents hydrogen bond donors outward from the center is bipolarized by hydrogen bond donors in the center of the pore and so prevent any proton leakage between water molecules (*13*). The Fab bound structures we report here do not impinge on the water channel or access of water to the channels, thus do not interfere with function. Multiple published studies have demonstrated that IgG binding does not impair water conductance of AQP4 (*14, 21, 22*). The extracellular surface epitopes are the structured extensions into the extracellular space. Antibody loops add further distance upon binding to the AQP4 epitopes from any access to the water channel. A combination of this, and the steric occlusion between M1 tetramer specific Fabs, and the interstitial binding of M23 favoring Fabs make it unlikely that any NMO antibodies would impede water channel function by binding. AQP4 internalization is likely the operative mechanism contributing to early myelin edema in NMO lesions (*23*).

The interaction of autoimmune antibodies with their target epitopes raises the question as to whether the antibodies modify the epitopes or are dominated by binding the intact configuration of the target epitopes. In the two cases we describe here, the epitope is remarkably constant in its configuration before and after Fab binding. This suggests that antibodies are not only selected to bind to target epitopes without plastic adaptation of the epitope but do so in a manner that compensates for the thermodynamic consequences of removing epitope from solvation by balancing for the entropic gain of desolvation. To assess the mode of AQP4 rAb binding, their impact on the conformation of the extracellular loops, and the molecular basis of their affinity and epitope specificity, we determined the atomic structures of two AQP4 rAbs derived Fab fragments bound to M1 AQP4, and extended the analysis to neighboring tetramers in M23 OAPs (*11*). The structures reveal why one of the antibodies rAb186 binds more tightly to OAPs than to AQP4 tetramers (*16*) (Fig 4, 5).

Loops A, C, E form highly ordered stable structures within the tetramer that are surprisingly constant and unaltered by bound Fabs. This accounts for the lack of Fab binding to linear peptides of the AQP4 loop epitopes alone since peptides would not conform to the structured conformation displayed on APQ4 (*10, 11*). In addition, identical loops from three or four different monomers contribute different orientations of the same structured sequences to the epitope resulting in a more complex assembly than for a tetramer alone. Hence, a sensitive diagnostic peptide binding assay to detect serum AQP4-IgG would need to retain some of the ordered displayed epitope. Furthermore, AQP4 OAP formation adds to epitope complexity, as the extracellular loops on multiple AQP4 tetramers are presented in close proximity.

The lattices of AQP4 are formed from protein-protein contacts that are completely different interfaces than the lipid mediated interfaces in AQP0, the only other case of aquaporin arrays (*24-26*). The unit cell dimension in AQP4 arrays is a=69Å (*17*) versus a=65.5Å in AQP0. Thus, the area of the unit cell in AQP4 corresponds to ∼8 more lipids per tetramer. In AQP4 OAPs, the tetramers are rotated to form true protein-protein contacts between tetramers opening a larger lipid filled area between four tetramers (Fig S11). While there is recognized physiological rationale for AQP0 lattices, the AQP4 lattices are structurally unrelated, and have not yet any recognized physiological reason for their presence.

Given the prominent role of AQP4-IgG in NMO lesion pathogenesis (*1-3*), direct inhibition of antibody binding would minimize damage and neurologic impairment. Interaction between AQP4-IgG and AQP4 may modulate both antibody effector function and the local inflammatory milieu, thereby modifying glial and neuronal injury (*15, 27*). The 3D structure of rAbs and AQP4-epitopes define interfaces that lead to assembly of antibody platforms that facilitate CDC. AQP4 rAbs sensitive to C loop mutations at H151/L154 enhance complement C1q activation on AQP4 OAPs by facilitating assembly of hexameric antibody complexes on AQP4 OAPs (*15, 28*). The two atomic structures of Fab bound AQP4 provide a template that instructs the design of small molecule or peptide inhibitors that block serum AQP4-IgG binding by directly competing with antibody binding or altering the structure of the epitopes on AQP4. Previous attempts to select inhibitory compounds have not been informed by a blueprint of the direct interactions between AQP4-IgG and AQP4 (*29*). These structures demonstrate the intricacy of composite epitope recognition and provide a platform for intelligent inhibitor design.

---

**Table 1.**
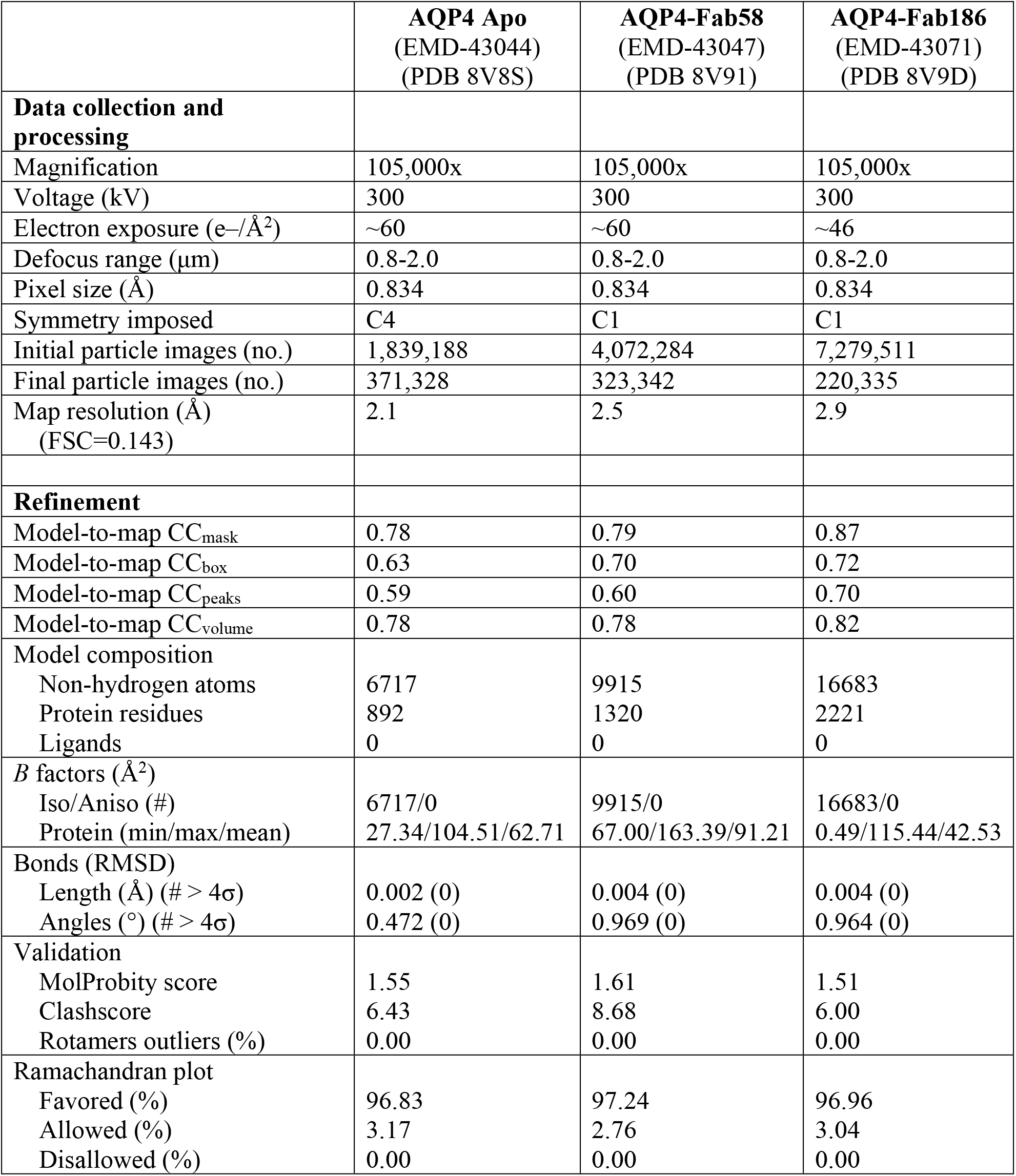
CryoEM data collection and processing.

|  | **AQP4 Apo**<br>(EMD-43044)<br>(PDB 8V8S) | **AQP4-Fab58**<br>(EMD-43047)<br>(PDB 8V91) | **AQP4-Fab186**<br>(EMD-43071)<br>(PDB 8V9D) |
| --- | --- | --- | --- |
| **Data collection and processing** |  |  |  |
| Magnification | 105,000x | 105,000x | 105,000x |
| Voltage (kV) | 300 | 300 | 300 |
| Electron exposure (e-/Å<sup>2</sup>) | ~60 | ~60 | ~46 |
| Defocus range (μm) | 0.8-2.0 | 0.8-2.0 | 0.8-2.0 |
| Pixel size (Å) | 0.834 | 0.834 | 0.834 |
| Symmetry imposed | C4 | C1 | C1 |
| Initial particle images (no.) | 1,839,188 | 4,072,284 | 7,279,511 |
| Final particle images (no.) | 371,328 | 323,342 | 220,335 |
| Map resolution (Å)<br>(FSC=0.143) | 2.1 | 2.5 | 2.9 |
| **Refinement** |  |  |  |
| Model-to-map CC<sub>mask</sub> | 0.78 | 0.79 | 0.87 |
| Model-to-map CC<sub>box</sub> | 0.63 | 0.70 | 0.72 |
| Model-to-map CC<sub>peaks</sub> | 0.59 | 0.60 | 0.70 |
| Model-to-map CC<sub>volume</sub> | 0.78 | 0.78 | 0.82 |
| Model composition |  |  |  |
| Non-hydrogen atoms | 6717 | 9915 | 16683 |
| Protein residues | 892 | 1320 | 2221 |
| Ligands | 0 | 0 | 0 |
| *B* factors (Å<sup>2</sup>) |  |  |  |
| Iso/Aniso (#) | 6717/0 | 9915/0 | 16683/0 |
| Protein (min/max/mean) | 27.34/104.51/62.71 | 67.00/163.39/91.21 | 0.49/115.44/42.53 |
| Bonds (RMSD) |  |  |  |
| Length (Å) (# > 4σ) | 0.002 (0) | 0.004 (0) | 0.004 (0) |
| Angles (°) (# > 4σ) | 0.472 (0) | 0.969 (0) | 0.964 (0) |
| Validation |  |  |  |
| MolProbity score | 1.55 | 1.61 | 1.51 |
| Clashscore | 6.43 | 8.68 | 6.00 |
| Rotamers outliers (%) | 0.00 | 0.00 | 0.00 |
| Ramachandran plot |  |  |  |
| Favored (%) | 96.83 | 97.24 | 96.96 |
| Allowed (%) | 3.17 | 2.76 | 3.04 |
| Disallowed (%) | 0.00 | 0.00 | 0.00 |

## Acknowledgments

We thank Laura Caboni, Daniel Asarnow, Miles Sasha Dickinson, Yifan Cheng who helped with initial discussions. Janet Finer-Moore read the manuscript and provided valuable input. David Bulkley, Glenn Gilbert, Matt Harrington supported with management of UCSF cryoEM facility for data collection and storage.

## Author contributions

Conceptualization: MG, RMS

Methodology: MG, NKK, AN, PH, SP, JLB, RMS

Investigation: MG, NKK, AN, PH, SP, JLB, RMS

Visualization: MG, NKK, RMS

Funding acquisition: MG, RMS

Project administration: MG, RMS

Supervision: MG, JLB, RMS

Writing – original draft: MG, NKK, RMS

Writing – review & editing: MG, NKK, JLB, RMS

## Data and materials availability

The models and maps obtained using cryoEM are deposited to the public repositories PDB and EMDB. The accession numbers to these databases are-AQP4 Apo in nanodiscs (PDB: 8V8S; EMDB: EMD-43044), AQP4 with Fab58 (PDB: 8V91; EMDB: EMD-43047), AQP4 with Fab186 (PDB: 8V9D; EMDB: EMD-43071). Other materials associated with this manuscript like plasmid used for expression of AQP4, are available upon request. Detailed protocols are provided in the supplementary materials.

