## Supplementary Materials for "Structural Basis of Aquaporin-4 Autoantibody Binding in Neuromyelitis Optica"

for

[1]Department of Biochemistry and Biophysics, University of California San Francisco; San Francisco, CA 94143, USA.
[2]current address: Department of Chemical Physiology and Biochemistry, Oregon Health & Science University, Portland, OR, 97239, USA.
[3]Departments of Neurology and Ophthalmology, Programs in Neuroscience and Immunology, University of Colorado School of Medicine, Anschutz Medical Campus; Aurora, CO 80045, USA.

**Supplementary Figure 1**

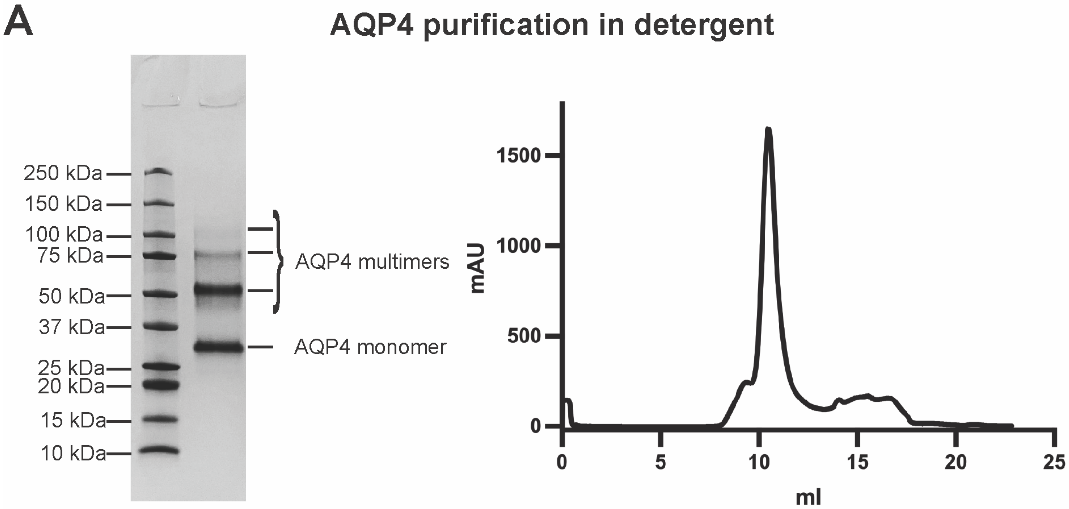


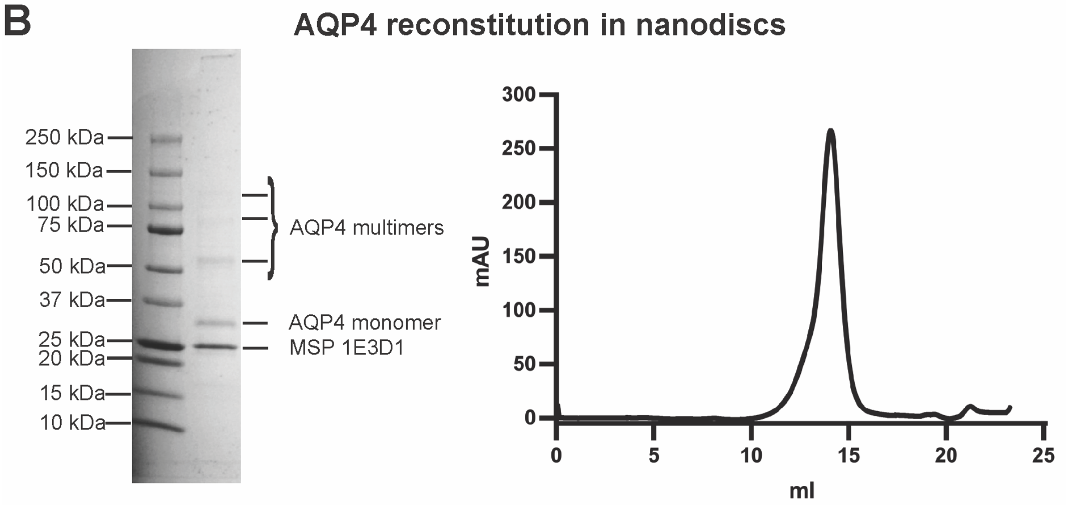


**Figure S1. Human AQP4 purification and nanodisc reconstitution.** **(A)** Purified AQP4 in detergent analyzed on SDS-PAGE and SEC profile of AQP4 in detergent using Superdex™ 200 Increase 10/300 GL column. **(B)** Reconstitution of purified AQP4 in lipid nanodiscs- SDS-PAGE and SEC profile using Superose™ 6 Increase 10/300 GL column.

**Supplementary Figure 2**

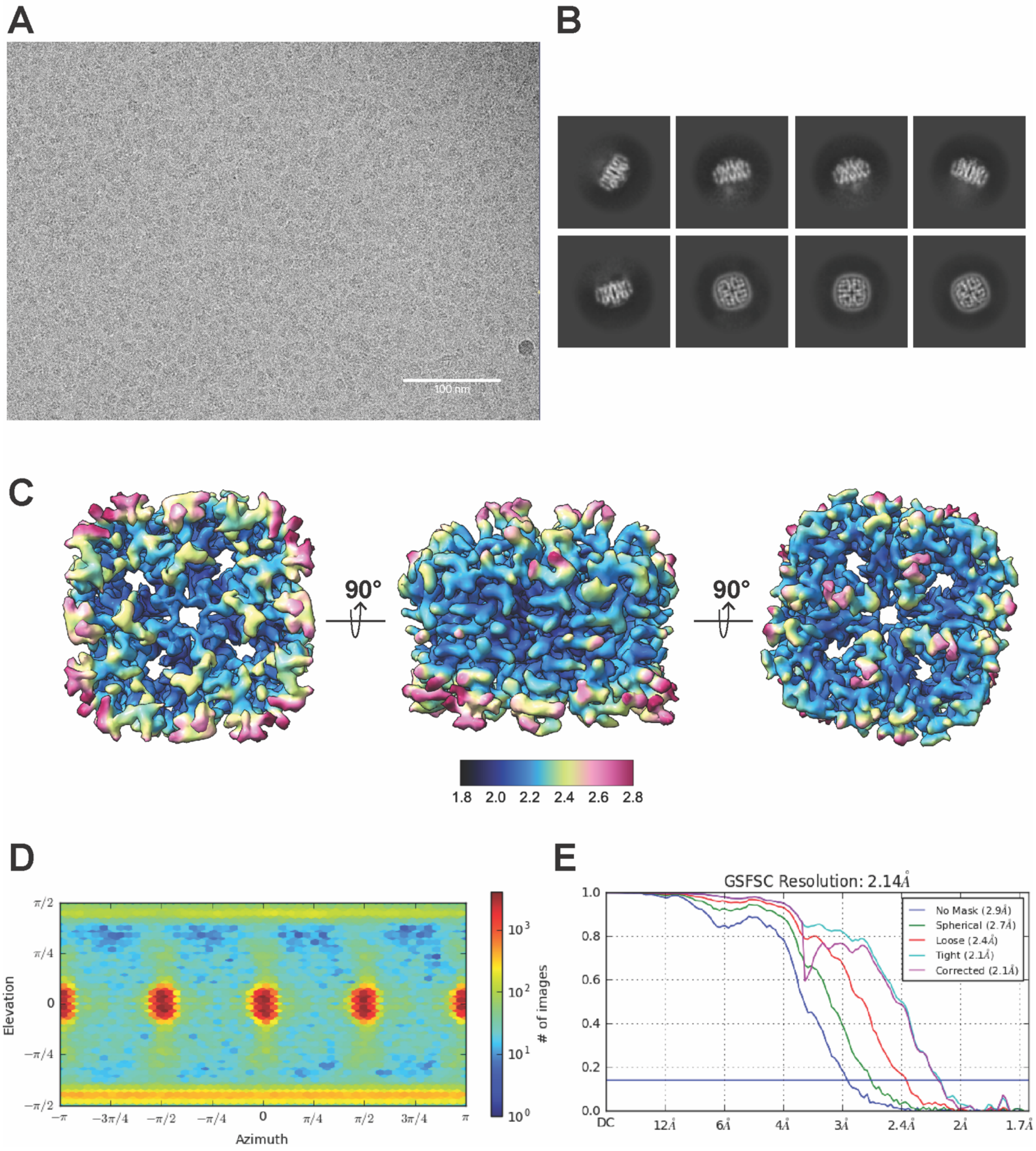


**Figure S2. AQP4 apo cryoEM data collection and processing. (A)** Representative micrograph from cryoEM experiment after motion correction. **(B)** Representative 2D classes from cryoEM data processing. **(C)** The top, side and bottom view of the final map obtained from the cryoEM data processing. **(D)** Euler angle distribution of the particle images.

**Supplementary Figure 3**

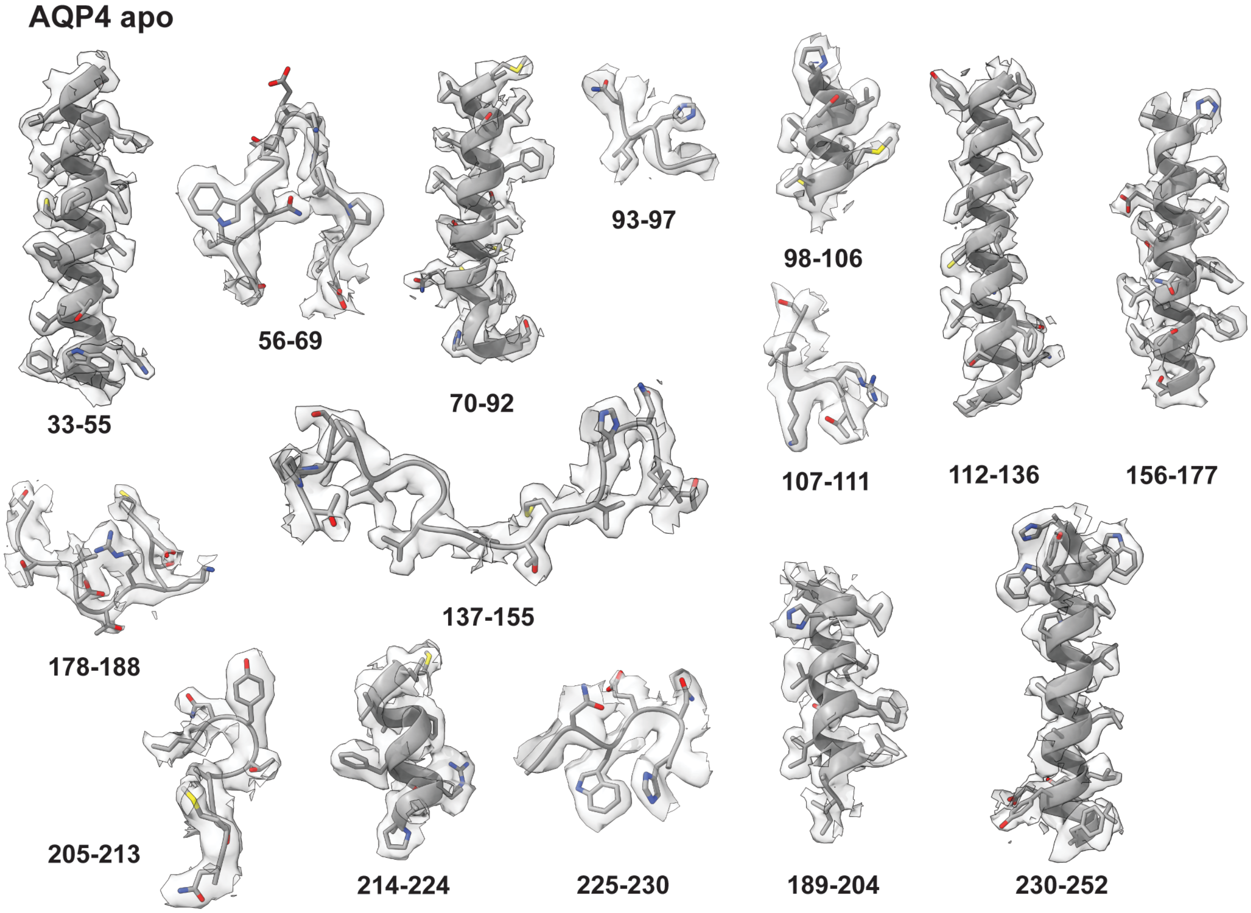


**Figure S3. AQP4 apo representative densities.** EM densities for the transmembrane helices and loops for the AQP4 apo with the model built into it.

**Supplementary Figure 4**

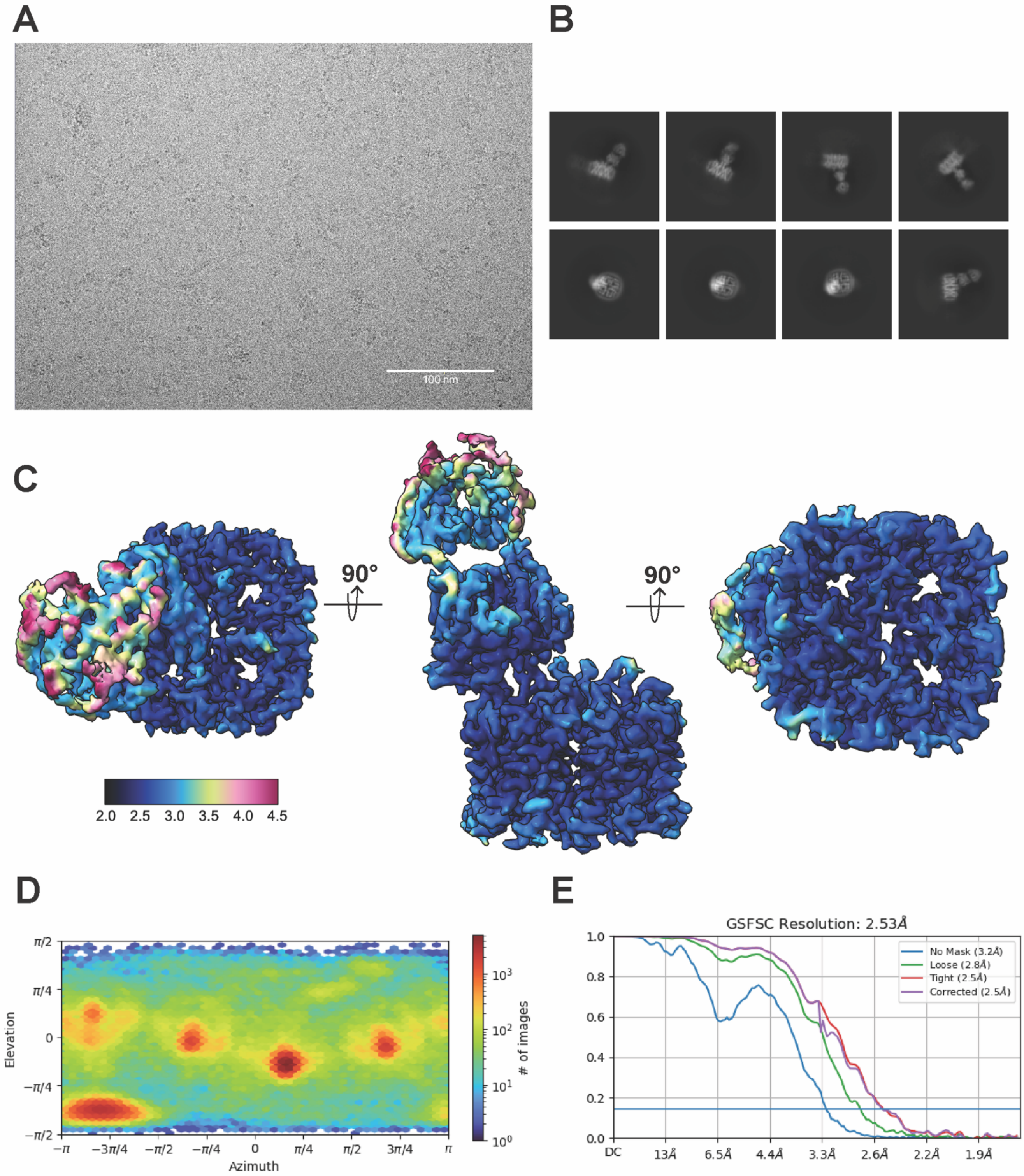


**Figure S4. AQP4-Fab58 cryoEM data collection and processing. (A)** Representative micrograph from cryoEM experiment after motion correction. **(B)** Representative 2D classes from cryoEM data processing. **(C)** The top, side and bottom view of the final map obtained from the cryoEM data processing. **(D)** Euler angle distribution of the particle images.

**Supplementary Figure 5**

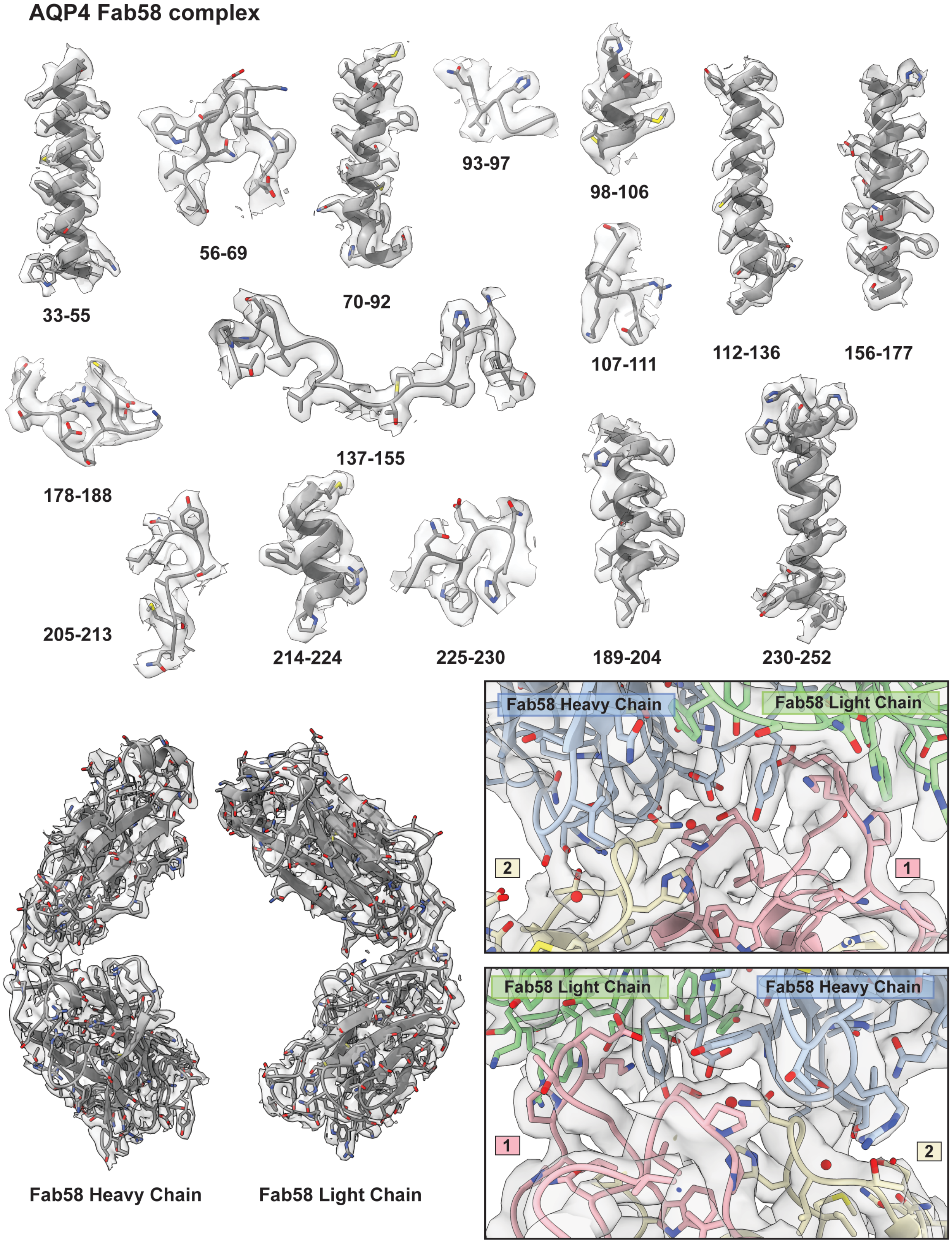


**Figure S5. AQP4-Fab58 representative densities.** EM densities for the transmembrane helices and loops for the AQP4-Fab58 with the model built into it.

**Supplementary Figure 6**

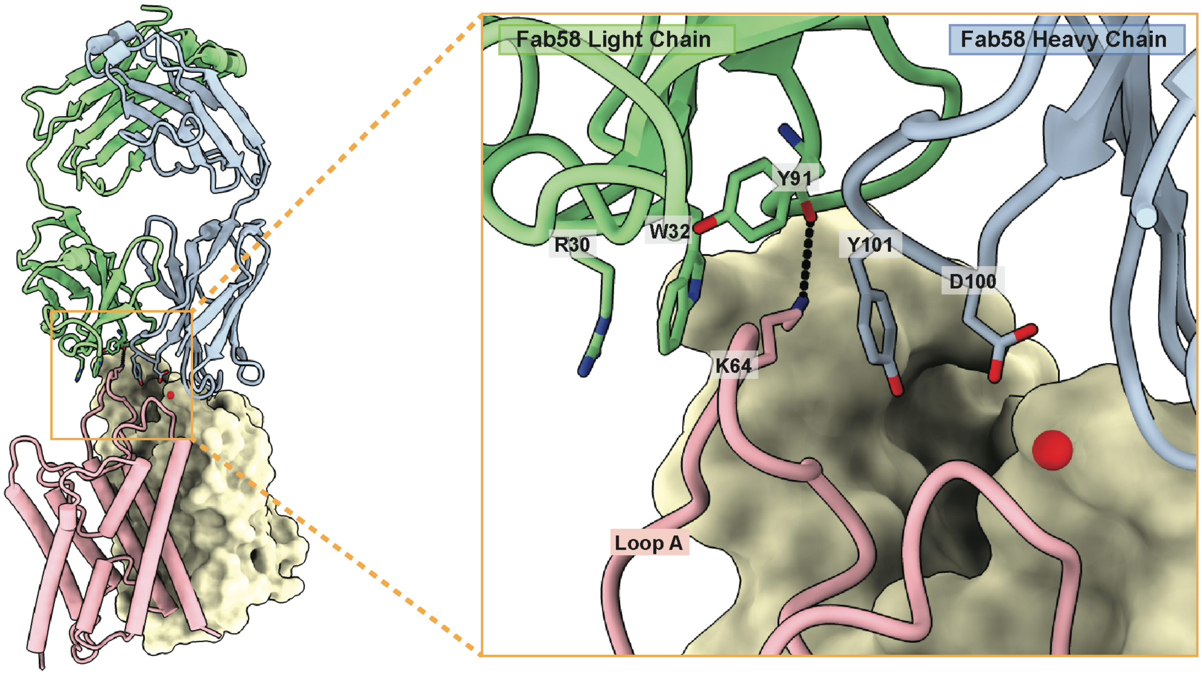


**Figure S6. AQP4-Fab58 loop A interactions.** Closer look of loop A interactions with HC and LC of Fab58.

**Supplementary Figure 7**

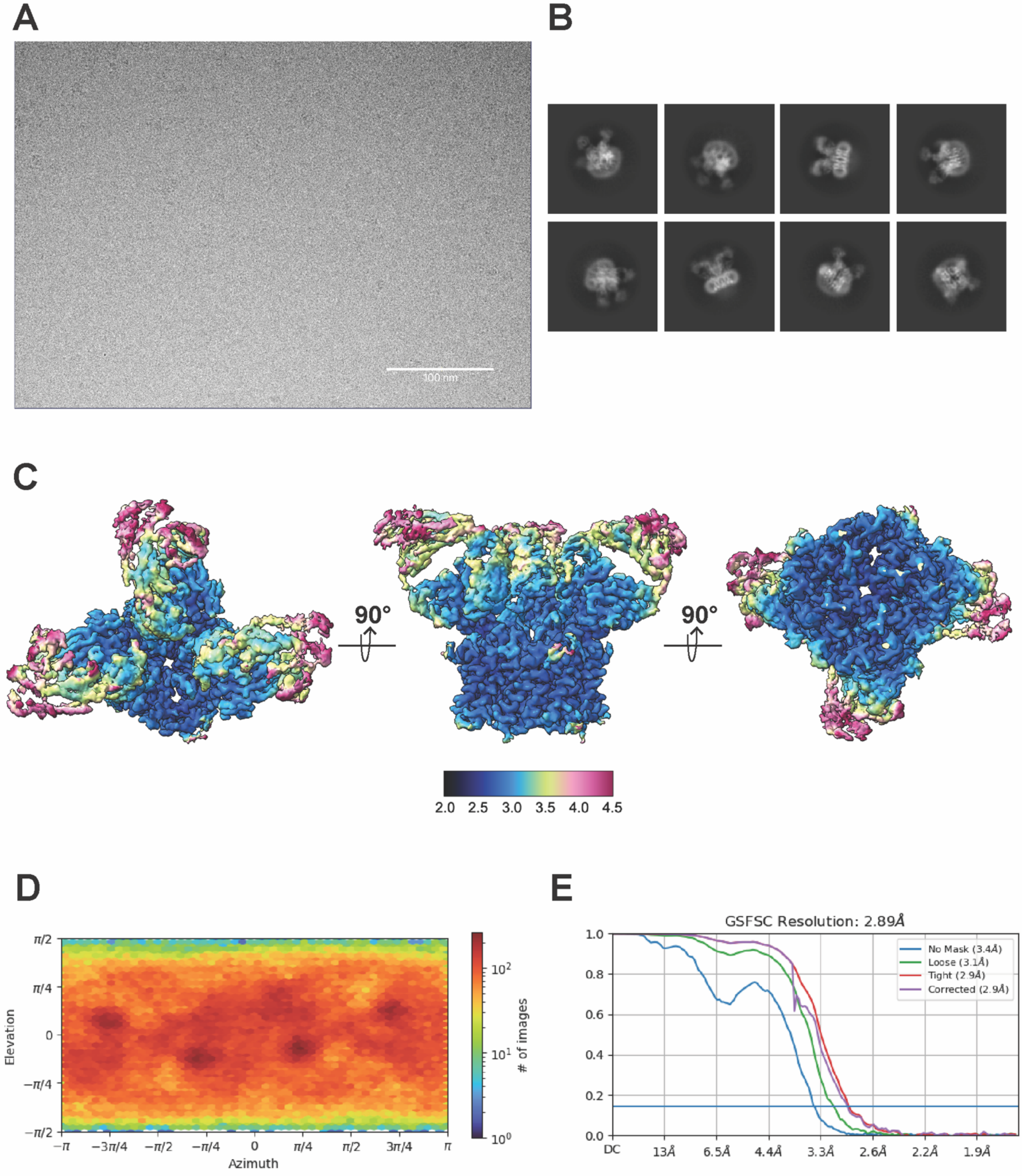


**Figure S7. AQP4-Fab186 cryoEM data collection and processing. (A)** Representative micrograph from cryoEM experiment after motion correction. **(B)** Representative 2D classes from cryoEM data processing. **(C)** The top, side and bottom view of the final map obtained from the cryoEM data processing. **(D)** Euler angle distribution of the particle images.

**Supplementary Figure 8**

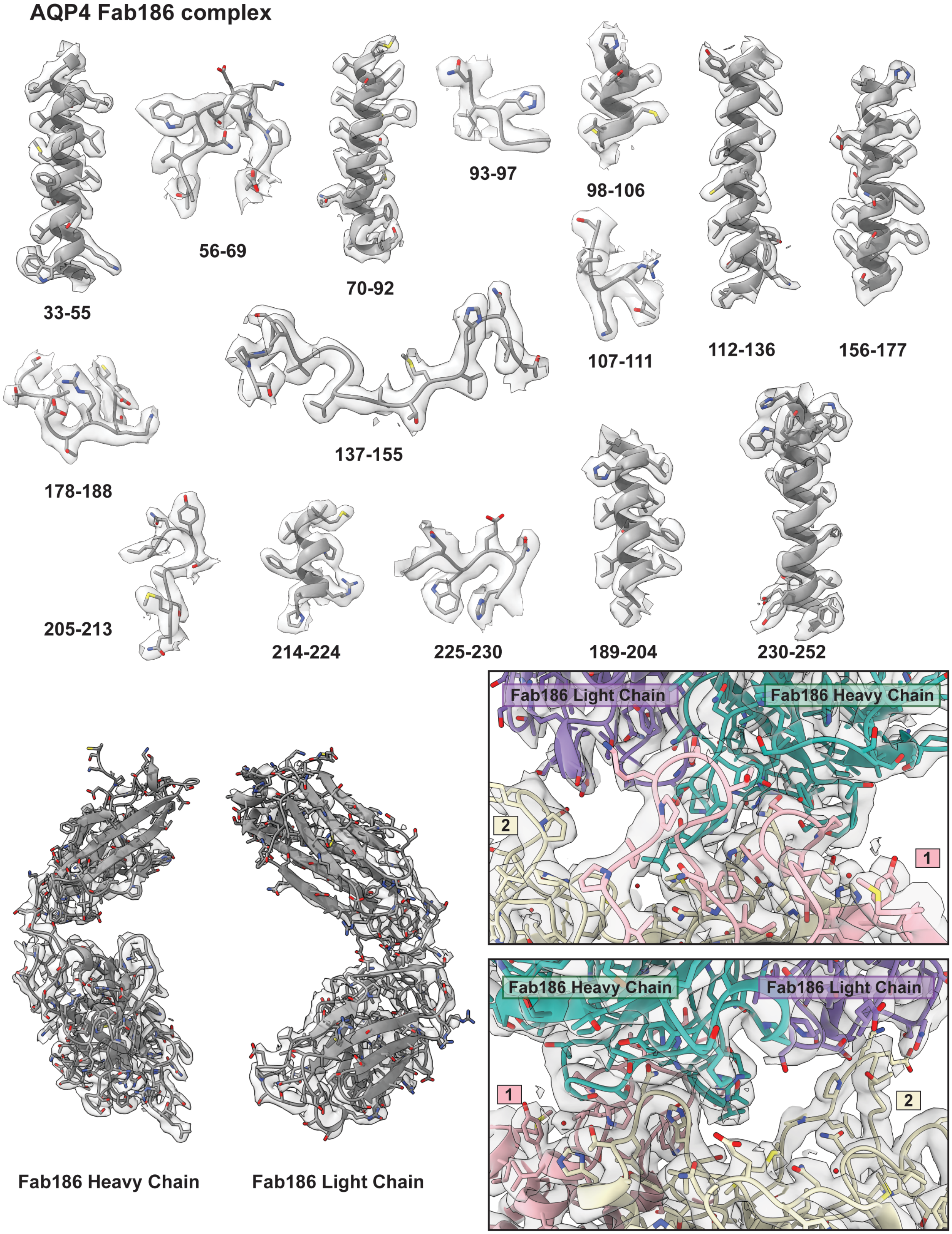


**Figure S8. AQP4-Fab186 representative densities.** EM densities for the transmembrane helices and loops for the AQP4-Fab186 with the model built into it.

**Supplementary Figure 9**

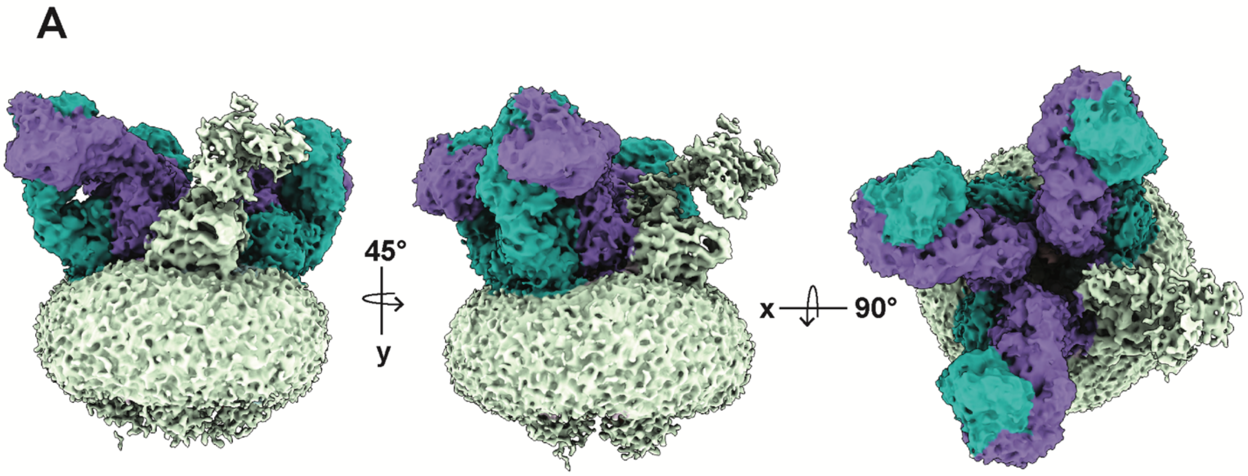


**Figure S9. AQP4-Fab186 low contour map exhibiting the density for a fourth Fab186.** The final cryoEM map shows densities prominently for three Fab186 bound to the but there is a weak density for the fourth Fab molecule. The fourth density is clearly visible in the 2D classes as well.

**Supplementary Figure 10**

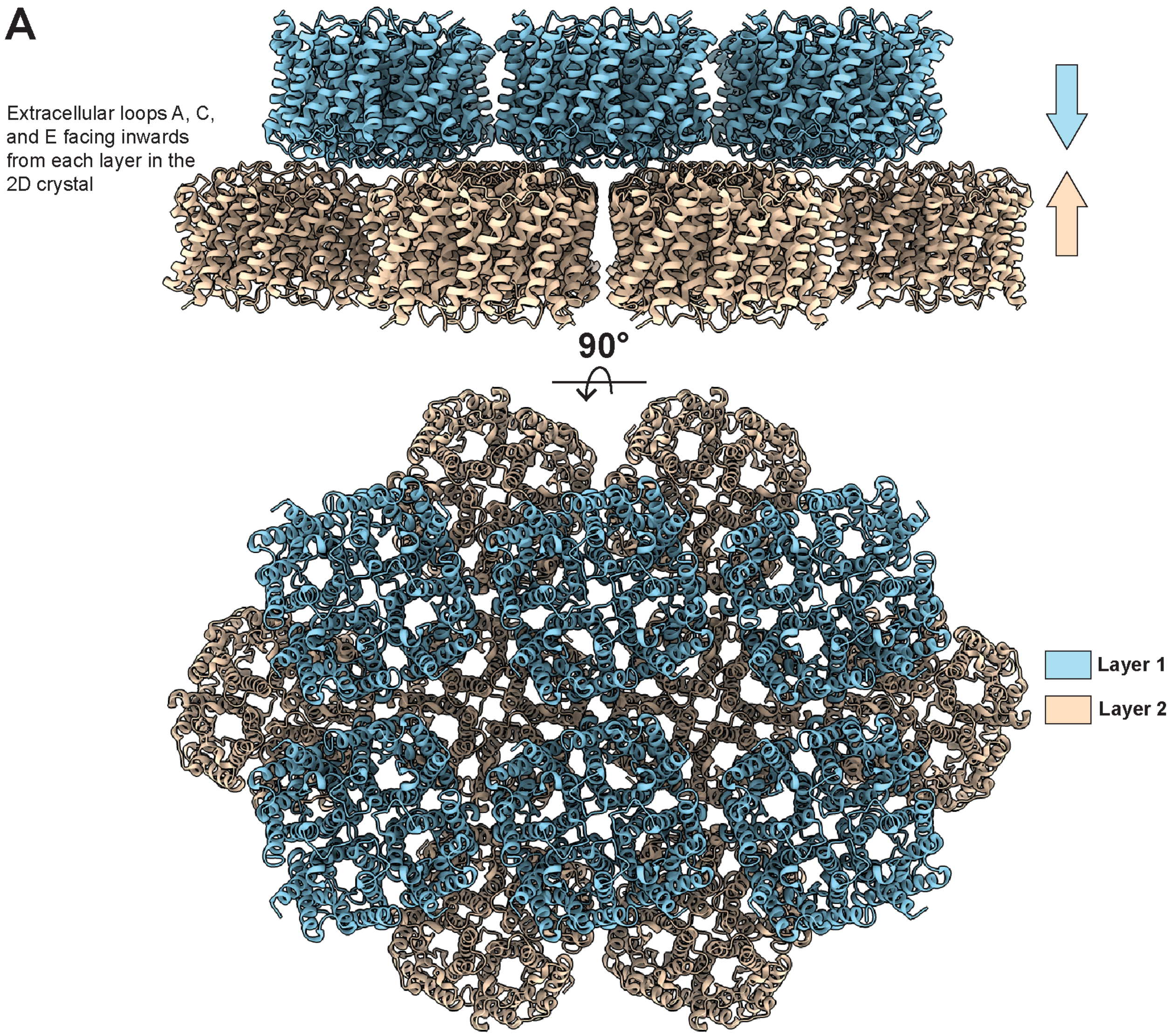


**Figure S10. Representation of two AQP4 layers in the 2D crystal. (A)** Side view and top view of the 2D crystal structure of rat AQP4 M23 isoform (PDB ID 2D57) exhibiting two layers of AQP4 array with the extracellular loops facing inwards. Layer 1 in color sky blue and Layer 2 in color peach.

**Supplementary Figure 11**

A

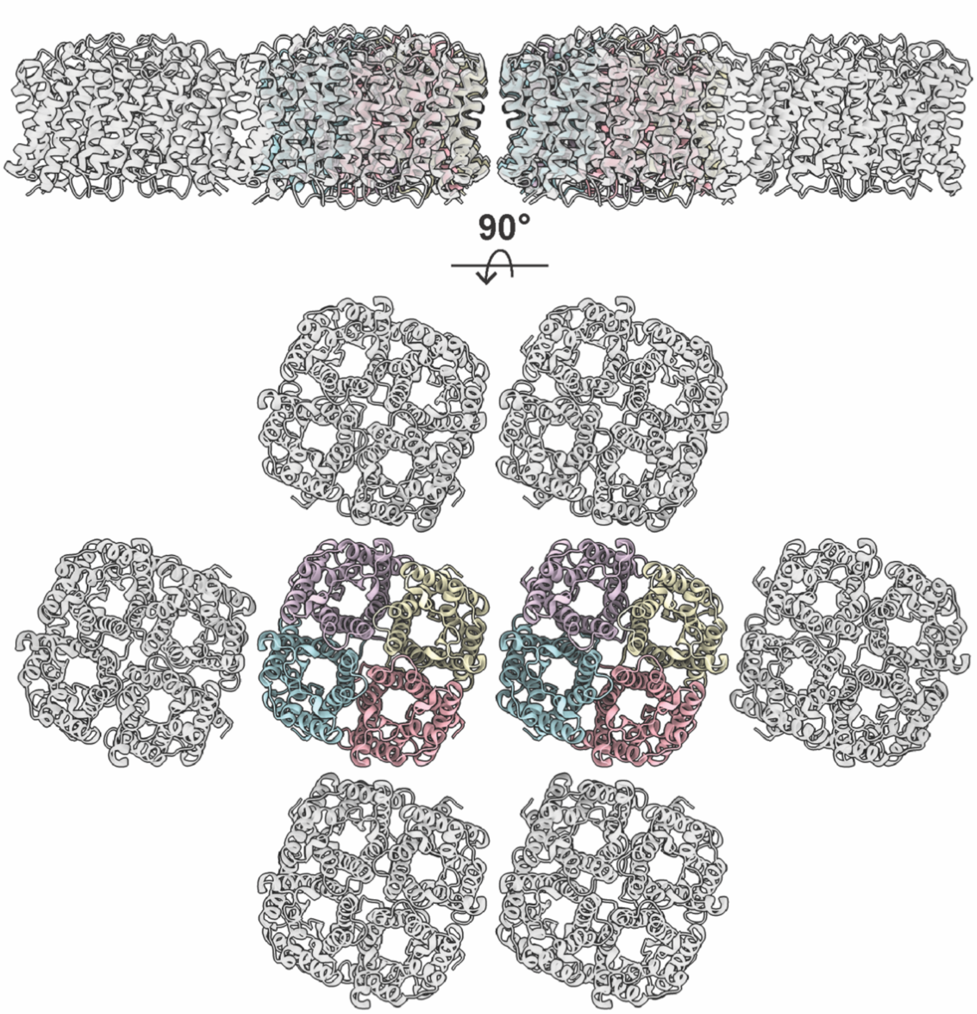


B

```
AQP4_HUMAN      MSDRPTARRWGKCGPLCTRENIMVAFKGVWTQAFWKAVTAEFLAMLIFVLLSLGSTINWG    60
AQP4_RAT        MSDGAAARRWGKCGPPCSRESIMVAFKGVWTQAFWKAVTAEFLAMLIFVLLSVGSTINWG    60
                ***   :********* *:**.******************************:*******

AQP4_HUMAN      GTEKPLPVDMVLISLCFGLSIATMVQCFGHISGGHINPAVTVAMVCTRKISIAKSVFYIA    120
AQP4_RAT        GSENPLPVDMVLISLCFGLSIATMVQCFGHISGGHINPAVTVAMVCTRKISIAKSVFYIT    120
                *:*:*******************************************************:

AQP4_HUMAN      AQCLGAIIGAGILYLVTPPSVVGGLGVTMVHGNLTAGHGLLVELIITFQLVFTIFASCDS    180
AQP4_RAT        AQCLGAIIGAGILYLVTPPSVVGGLGVTTVHGNLTAGHGLLVELIITFQLVFTIFASCDS    180
                **************************** *******************************

AQP4_HUMAN      KRTDVTGSIALAIGFSVAIGHLFAINYTGASMNPARSFGPAVIMGNWENHWIYWVGPIIG    240
AQP4_RAT        KRTDVTGSVALAIGFSVAIGHLFAINYTGASMNPARSFGPAVIMGNWENHWIYWVGPIIG    240
                ********:***************************************************

AQP4_HUMAN      AVLAGGLYEYVFCPDVEFKRRFKEAFSKAAQQTKGSYMEVEDNRSQVETDDLILKPGVVH    300
AQP4_RAT        AVLAGALYEYVFCPDVELKRRLKEAFSKAAQQTKGSYMEVEDNRSQVETEDLILKPGVVH    300
                *****.***********:***:***************************:**********

AQP4_HUMAN      VIDVDRGEEKKGKDQSGEVLSSV 323
AQP4_RAT        VIDIDRGDEKKGKDSSGEVLSSV 323
                ***:***:******.********
```

**Figure S11. Estimation of spatial arrangement of AQP4 in OAPs. (A)** Side view and top view of the single layer of AQP4 array derived from 2D crystal. For the two representative neighboring tetramers same color scheme as in other figures was used. **(B)** Sequence comparison of Human and Rat AQP4.

**Movie S1**
**Morph between Fab58 and Fab186 on two AQP4 monomers.** Both Fab58 and Fab186 interactions span over two monomers within a AQP4 tetramer. To make a comparison of their interaction with respect to AQP4, this linear morph represents transition from Fab58 binding to Fab186 binding over two monomers.

**Movie S2**
**Spin movie showing potential rAB186 association with OAPs.** This is a spin movie for the images shown in Figure 5 to emphasize on the proximity of lateral surface of Fab186 to the extracellular loops of the neighboring tetramer in an estimated OAP.

### Materials and Methods

#### AQP4 expression and purification

The full-length human aquaporin 4 gene isoform M1 (AQP4 M1) was synthesized and cloned in pPICZ expression vector (Genscript). The expression construct was designed with an N-terminal 8xHis followed by a flag tag (DYKDDDDK) and a thrombin cleavage site and cloned into the EcoR1 and Not1 sites of the vector. A protocol similar to previously published was used for the expression of AQP4 M1 in *Pichia pastoris* with modifications (*13*). In summary, the expression vector was electroporated into *P. pastoris* X-33 cells and the transformed cells were then selected as colonies on YPD plates with 50 g/mL Zeocin (Invitrogen) and tested for expression. More than 100 colonies were screened in batches and compared to the best AQP4 expressor of each round to yield a high-expression stable strain of *P. pastoris* (MG3). For protein production, the *P. pastoris* MG3 strain was cultured in growth media at 30°C for 24 h, then the temperature was lowered to 26°C, and methanol was added directly to the cultures to a final concentration of 2.5%. The cultures were grown for another 48 h. Cultures were harvested by centrifugation at 4°C, 6,000*xg* for 10 min. Cells were resuspended in the lysis buffer (1X TBS with 2mM βME) and lysed by bead beating with glass beads. Broken and unlysed cells were removed by centrifugation at 4°C, 6,000*xg* for 10 min. The supernatant was pelleted at 160,000*xg* at 4°C for 2 h to extract the membranes. Pellets were resuspended in MR Buffer (25 mM Tris HCl, pH 7.4 at room temperature, 250 mM NaCl, 10% glycerol, 1 mM βME) and stored at -80°C until further processing. To begin purification, resuspended membranes were solubilized by adding 1% n-dodecyl-β-D-maltoside (DDM) (Anatrace) to a final concentration of 200 mM and stirred at 4°C for 1 h. Unsolubilized material was pelleted at 160,000*xg* at 4°C for 30 min. 50 mM imidazole, pH 7.4, was added to the supernatant. The supernatant was then batch bound with Ni-NTA resins (Qiagen) for 2 h, loaded onto a Bio-Rad Econo Column and washed with MR Buffer with 30 mM βDDM and 50 mM imidazole, and then eluted with 300 mM imidazole. Imidazole was removed using Econo-Pac DG10 desalting column (Bio-Rad) equilibrated with MR Buffer with 30 mM βDDM. The N-terminal tag was cleaved by thrombin at 4°C overnight. Uncleaved AQP4 was removed the next day with HisPur™ Cobalt Resin (Thermo Scientific). Then, AQP4 was concentrated in a 50,000 molecular weight cut-off Amicon spin concentrator (Millipore) and further purified by size exclusion chromatography on a Superdex™ 200 Increase 10/300 GL column (Cytiva) in 25mM Citrate, pH 6.0, 50 mM NaCl, 5% glycerol, 30 mM βDDM, and 2 mM DTT.

#### AQP4 nanodisc reconstitution

Purified AQP4 in detergent with the intact affinity tags was used for this reconstitution. 100 μM of AQP4 was mixed with 300 μM of purified 1E3D1 membrane scaffold protein (MSP), and 10 mM of Soy Polar Lipids (Avanti). The mixture was incubated overnight at 4°C with Bio-Beads® SM2 resin (Bio-Rad) to adsorb the detergent from the solution. The sample was separated from Bio-beads and allowed to bind to HisPur™ Cobalt Resin (Thermo Scientific) to separate reconstituted nanodiscs with AQP4 and remove empty nanodiscs using affinity tag on AQP4. Reconstituted AQP4 in nanodiscs was then eluted with 150 mM imidazole containing buffer. The elute was concentrated using 50,000 kDa molecular weight cut-off Amicon spin concentrator (Millipore) and further purified by size exclusion chromatography on a Superose™ 6 Increase 10/300 GL column (Cytiva) in 50 mM HEPES, pH 7.5, 100 mM NaCl buffer.

#### rAb expression and purification

Recombinant monoclonal NMO antibodies rAb58 and rAb186 were generated from clonally expanded cerebrospinal fluid plasma blasts as described (*5, 16*). VH and VL constructs were co-transfected (1:3 ratio) into Expi293F cells for expression. The supernatant was harvested, centrifuged to remove any cells and debris, adjusted to pH 8, and purified at 4°C on an AKTA pure 25 system using a HiTrap MabSelect affinity column and HiPrep 26/10 desalting column (Cytiva). The rAb was eluted from the MabSelect column in 0.1 M glycine (pH 2.7) and desalted using 1x phosphate buffered saline (PBS). Desalted rAb was adjusted to pH 7.5 with 1 M Tris-HCl, pH 8.0 and exchanged and concentrated in PBS using Ultracel YM-30 micro concentrators (Millipore, Billerica, MA). Antibody integrity and Fab fragments were confirmed by denaturing and native-PAGE, and IgG concentration was assayed using a Nanodrop spectrophotometer (Thermo Fisher Scientific).

**Fab preparation**
Pierce™ Fab purification kit (Thermo Scientific) was used to make Fabs from the IgGs following the manufacturer's standard protocol. In summary, papain protease cleavage was performed to digest human IgGs resulting in Fab and Fc fragments. The Fc fragment was separated using Protein A agarose. The purity of Fab was assessed on SDS-PAGE gel.

**Bio-Layer Interferometry**
The MSP 1E3D1 was biotinylated before AQP4 nanodisc reconstitution. Then AQP4 in biotinylated MSP 1E3D1 was immobilized on Octet® Bio-Layer Interferometry (BLI) Streptavidin (SA) Biosensor tips (Sartorius). Patient-derived recombinant IgGs were kept in solution at various concentrations. Octet Red 384 system was used at 25°C with 1000 rpm shaking for the measurements. The IgG binding was assayed using BLI on the immobilized AQP4 tips using a 16-channel experiment. Dissociation coefficients ($K_d$) were estimated assuming heterogenous ligand fitting model. An IgG isotype control was used to assess the baseline of the experiment.

**CryoEM sample preparation**
AQP4 M1 isoform in 1E3D1 nanodiscs were used at 0.3 μM for the apo structure. For AQP4-Fab58 sample, AQP4 in nanodiscs was mixed with ~5-fold molar excess of Fab58. AQP4-Fab186 sample was prepared in a similar way. Final concentration of AQP4 in the mixture was 0.3 μM at the time of grid freezing in all the samples. A mixed population of cryoEM 2D-classes of 1-4 four Fab186 bound AQP4 tetramers was obtained hence an increased the Fab186 ratio to 30-fold molar excess that of AQP4 was used. This resulted in saturation of all four binding sites on the AQP4 tetramer. There was a severe orientation bias with this sample that we were not able to rectify even after employing multiple data collection and processing strategies. This prohibited us from achieving a high-resolution map. AQP4 in detergent was used for obtaining the Fab186 bound structure where 18 μM of AQP4 was mixed with ~75 μM of Fab186. Final AQP4 (in detergent) concentration on grids was ~12 μM.
For grid preparation, Mark IV Vitrobot (FEI) was used at 10°C and 100% humidity. 3 μL of the sample was applied to freshly glow-discharged Quantifoil R 1.2/1.3 400-mesh Au holey carbon grids. Blotting time used was between 3-7 s with a blot force of -2 to remove excess sample from the grids. Rapid plunge freezing was done in liquid ethane after blotting and grids were stored in liquid nitrogen.

**CryoEM data collection**
CryoEM data acquisition was on Titan Krios, a 300kV transmission electron microscope, at a defocus range from -0.8 to -2.5, a total dose between 45-60 e/Å$^2$. The Titan Krios was operated with a post-column energy filter (20 eV slit width) and GATAN K3 direct electron detector. The data collection was setup using Serial-EM with automated data acquisition features. Number of micrographs collected for each structure, dose, pixel size, are listed in supplementary table 1. For AQP4-Fab58 structure, two datasets- one at 0° stage tilt and another at 30° stage tilt was collected. Addition of 30° tilt data helped us overcome the orientation bias problem. For AQP4-Fab186 structure, the datasets were collected at 0° stage tilt and 20° stage tilt for the same reason.

**CryoEM data processing and model building**

The movies obtained from data collection was subjected to dose-weighted beam-induced motion correction using MotionCor2 to obtain the micrographs. CryoSPARC (v3.2 and v4.2.1) was used for further data processing (*30*). Briefly, the Patch CTF in CryoSPARC was used for CTF estimation and the micrographs were manually curated. The 0° and 30° stage tilt data were merged for Fab58 after this step. Similarly, 0° and 20° stage tilt data were merged for Fab186 data. Blob picker was used for particle picking to generate templates for eventual template picking of the particles. After particles picking, 2D classification was performed. It was followed by ab-initio 3D volume generation and heterogenous classification using ab-initio volumes. Multiple rounds of 2D classification, ab-initio volume generation and heterogenous classification was performed. The best quality map from these steps was used for non-uniform refinement. C4 symmetry was imposed for the AQP4 apo structure while AQP4-Fab58 and AQP4-Fab186 structures were determined without any imposed symmetry during data processing.

For model building of AQP4 part of the structures, we used our previously published crystal structure model as the starting point (PDB 3GD8) (*13*). We used Coot for model building and refinements in Phenix, software hosted by SBGrid consortium (*31*). For the Fab fragments, AlphaFold model was first generated using the known amino acid sequences (*32*). The predicted model was then refined in Coot and Phenix according to the cryoEM map obtained.

**OAP estimation**

Hiroaki et. al. determined the structure of the rat M23 isoform of AQP4 that forms a double bilayer assembly in which the external faces of AQP4 tetramers in each layer associate to for a double bilayer. 3D-crystal structure was determined by electron diffraction to a resolution of 3.2Å in plane/3.6Å perpendicular to the plane of the double bilayers (*17*) (Fig S10). This association (PDB ID 2D57) resulted in deformation of the external loops as compared to our previously published crystal structure of AQP4 and cryoEM structures in this manuscript (*13*). The cell dimensions in OAPs are identical to those of the double layer, and may have a role, arguably though not proven in cell-cell adhesion. Hence, one layer of the double bilayer interface was used as the template for OAPs (Fig S10). We therefore superimposed our experimentally determined AQP4 tetramer structure into each unit in the array (Fig S11A). The rat and human AQP4 arrays have the same cell dimensions, and their sequences are highly conserved (Fig S11B).

To assess if Fab186 can potentially interact with a neighboring tetramer in an OAP, we used minimal unit of Fab186 interaction from our structures- one Fab186 molecule and two monomers from a tetramer. From our array estimation we extracted two adjacent tetramers. The minimal AQP4-Fab186 interaction unit was imposed on the two adjacent array tetramers (Fig 5).
